# Sweet taste receptor regulates proliferation and glucose uptake in glioblastoma cells

**DOI:** 10.1101/2025.06.27.661813

**Authors:** Ana R. Costa, Ana C. Duarte, Ana C. Jorge, Isabel Gonçalves, Carolina Lucas, Inês R. Matos, Cecília R.A. Santos

## Abstract

Glioblastoma is a brain tumour classified by the World Health Organization as grade 4 due to its aggressiveness, invasiveness, and poor differentiation. Current standard therapies remain largely ineffective, reinforcing the need for novel targets to improve glioblastoma prognosis. Reprogramming of cellular metabolism is an important hallmark of cancer, marked by a shift from oxidative phosphorylation to glycolysis as the primary energy source. This metabolic switch, known as the Warburg effect, is exacerbated by the reduced oxygen and glucose availability in the tumour microenvironment. The sweet taste receptor (STR) is an important glucosensor with recognised role in the regulation of glucose uptake in several organs and in astrocytes, the glioblastoma precursor cells. We hypothesised that STR may regulate glucose uptake and metabolism in glioblastoma, representing a potential anticancer target. To test this, we investigated the effects of STR inhibition with lactisole, a specific TAS1R3 inhibitor, in three glioblastoma cell lines (U-87MG, SNB-19 and U-373MG) and in the UPCI-SCC-154 tongue cancer cell line. We provided evidence that STR inhibition consistently reduced cell viability and migration, without inducing apoptosis or necrosis, and impaired glucose uptake and L-lactate production, particularly in tongue cancer UPCI-SCC-154, and glioblastoma U-87MG and SNB-19 cells. Moreover, in a 3D spheroid model, lactisole reduced the invasion capacity of U-87MG glioblastoma spheroids. These results suggest that STR contributes to glioblastoma cell metabolism and behaviour and may represent a promising therapeutic target.

## 1. Introduction

Glioblastoma is a highly aggressive astrocytoma classified by the World Health Organization as a grade 4 brain tumour (Louis et al. 2021). The current standard treatment for glioblastoma includes surgical resection followed by radiotherapy and temozolomide (TMZ), along with supportive medication to relieve neurological symptoms. Unfortunately, although this multimodal therapeutic approach is the best option available, it results in 2-year survival rates of 10–26% and 5-year survival rates of 0–10% after diagnosis (Ferlay et al. 2021; Reni et al. 2017). The limited efficacy of current treatments is further hindered by the poor permeability of the blood-brain barrier to TMZ (Da Ros et al. 2018). In this context, the search for more effective therapies is both timely and crucial, and novel targets must be identified to improve prognosis.

An important hallmark of cancer, including glioblastoma, is the specificity of its energetic metabolism (Hanahan & Weinberg 2011; Pavlova & Thompson 2016; Taylor et al. 2019). Some cancer cells preferentially rely on glycolysis over oxidative phosphorylation for energy production, even in the presence of oxygen. This metabolic switch, first observed by Otto Warburg, favours the large demand of nucleotides, amino acids, and lipids essential for cell proliferation, and is enhanced by the reduced oxygen and glucose availability in the tumour microenvironment (Liberti & Locasale 2016; Mohanti et al. 1996; Yuen et al. 2016). Under hypoxic conditions, the hypoxia-inducible factor HIF1α is activated, which in turn transactivates several genes involved in glucose uptake, like glucose transporters, and glucose breakdown, such as hexokinase, phosphofructokinase, and aldolase, stimulating glycolysis and angiogenesis. Concurrently, glycolysis also releases lactate into the tumour microenvironment, which decreases the pH, weakening the immune response of the tumour surrounding cells and providing an additional energy source for tumour growth (Denko 2008). Moreover, glioblastoma chemoresistance might be enhanced by these metabolic features, as seen in other types of cancer (Desbats et al. 2020). Hence, the identification of metabolic targets to treat glioblastoma is promising. Reversing the Warburg effect in glioblastoma has been shown to reduce tumour size and aggressiveness, but only the glycolysis inhibitor 2-deoxyglucose is undergoing clinical trials to prevent the Warburg effect in glioblastoma (Raez et al. 2013; Sborov et al. 2015; Stein et al. 2010). Therefore, the identification of other potential players mediating the crosstalk between cell proliferation, invasion, hypoxia, and glucose metabolism in glioblastoma is of interest for the development of novel therapies targeting this aggressive tumour.

The taste receptor for sweet compounds (sweet taste receptor – STR) is an important glucosensor, with a recognised role in the regulation of glucose uptake in several organs, including the brain, by controlling the rate of glucose absorption and metabolism (Ren et al. 2009; Santos et al. 2019; Smith et al. 2018). The STR is composed by a heterodimer of TAS1R2 and TAS1R3 subunits that bind various sugars, artificial sweeteners, and sweet amino acids (Welcome & Mastorakis 2018). Although initially identified in the taste buds of the oral cavity, STR subunits are expressed throughout the body (Santos et al. 2019). Recent findings suggest that STR controls the glucose metabolism in astrocytes, the glioblastoma precursor cells, and in neurons of different nutrient-sensing forebrain regions (Ren et al. 2009; Smith et al. 2018). STR expression was also detected in cells in the third ventricle, which are sensitive to high concentrations of glucose (Ren et al. 2009; Welcome & Mastorakis 2018). This raises the hypothesis that STR might play a key role in regulating glucose uptake and metabolism in tumour cells, where energy demands are exceptionally high. Therefore, we hypothesize that STR expressed in glioblastoma cell lines and in human tumour samples of glioblastoma patients might function as sensor of the glucose availability in the tumour microenvironment. Our results show that STR inhibition with lactisole, a specific TAS1R3 inhibitor, reduced cell viability and migration, but did not induce apoptosis, in glioblastoma cell lines, particularly under oxygen and glucose deprivation, highlighting its potential as a therapeutic target in glioblastoma.

## 2. Materials and methods

### 2.1 Materials

For cell cultures, the Dulbecco’s Modified Eagle’s Medium (DMEM) with 4.5 g/L glucose (high glucose) with stable glutamine was purchased from bioWest (#L0103), and the Minimal Essential Medium was purchased from Sigma (#M0268).

MEM non-essential amino acids (NEAA; #11140035) and CellEvent^TM^ Caspase-3/7 Green Detection Reagent (#C10423) were purchased in ThermoFisher Scientific, and MTT [3-(4,5-dimethylthiazol-2-yl)-2,5-diphenyltetrazolium bromide] (#1006) was purchased in Gerbu Biotechnik GmbH.

Glucose Uptake Cell-Based (#600470) and L-Lactate (#700510) assay kits, and lactisole (CAS No 150436-68-3; #18657) were purchased from Cayman Chemical. A stock solution of lactisole 0.8 M was prepared in dimethyl sulfoxide (DMSO) and freshly dissolved in culture medium before the experiments. A vehicle control, where the DMSO final concentration did not exceed 0.625%, was included in all the experiments.

### 2.2 Cell Culture

The experimental procedures were performed using three commercial glioblastoma cell lines (U-87MG, SNB-19 and U-373MG), that present different characteristics and proliferation rates, in order the attempt to mimic glioblastoma heterogenicity. Human malignant glioblastoma cell lines grown in DMEM high glucose with stable glutamine supplemented with 10% (v/v) fetal bovine serum (FBS) and 0.1% (v/v) penicillin/streptomycin.

In addition, a tongue cancer cell line UPCI-SCC-154 was used as positive control in the experimental assays, since the taste receptors were firstly described in the oral cavity. The squamous carcinoma cells UPCI-SCC-154 were grown in MEM supplemented with 1x NEAA, 10% (v/v) FBS and 1% (v/v) penicillin/streptomycin.

All the cell lines were maintained under a controlled humidified atmosphere at 37°C and 5C% CO2.

### 2.3 Evaluation of the impact of TAS1R3 inhibition in tongue cancer and glioblastoma cells

#### 2.3.1 Effects of TAS1R3 inhibition in the cell proliferation

The cells’ viability was carried out using MTT assay. Briefly, tongue cancer UPCI-SCC-154, and U-87MG, SNB-19 and U-373MG glioblastoma cells were grown until 60% confluency and incubated for 48h at 37°C and 5% CO2 with control vehicle (DMSO 0.625%) or 5 mM lactisole, a known inhibitor of the sweet taste receptor TAS1R3 subunit. After 48 hours, 100 μL culture medium were removed and 10 μL of MTT solution (5 mg/mL in PBS) were added for approximately 45 min at 37°C in a humidified atmosphere containing 5% CO2. Ethanol 70% treated cells were used as positive control. Following MTT incubation, formazan crystals were dissolved in DMSO for 15 min, and absorbance was read at 570 nm in a microplate spectrophotometer xMark^TM^ (Bio-Rad). The cell viability was expressed as a percentage of the absorbance determined in the vehicle controls.

#### 2.3.2 Effects of TAS1R3 inhibition in the cell migration rate

Migration assays were performed to evaluate the cells capacity to migrate upon STR inhibition. Tongue cancer and glioblastoma cell lines were grown until 90-100% confluency, and then a scratch was created in each well by scraping a straight line with a micropipette tip. The debris were removed, and the edge of the scratch was smoothed by washing the cells once with PBS 1x. The appropriate culture medium was added to each well, in the presence or absence of 5 mM lactisole or DMSO (vehicle control), as described above. After that, each well was photographed at 0, 24 and 48 hours under an inverted microscope Axio Observer Z1 (Carl Zeiss) using a magnification of 5x (A-Plan 5x/0.12 Ph0 M27). For each image, the area of the scratch was measured using Fiji software (Schindelin et al. 2012), by individually comparing the images from the 24 and 48 hours timepoints to the measurements at the ground state (0 hours).

#### 2.3.3 Effects of TAS1R3 inhibition in the cell death

Cell apoptosis is a form of cell death, that unlike necrosis, is characterized by cell shrinkage, loss of membrane integrity and nuclear fragmentation without triggering inflammatory processes. Therefore, this assay was carried out to investigate whether inhibition of the sweet taste receptor TAS1R3 subunit with lactisole could deploy apoptosis or necrosis. To perform this assay, glioblastoma cells were stained with CellEvent^TM^ Caspase-3/7 Green kit to evaluate apoptosis and propidium iodide to evaluate cell death by necrosis. Briefly, the three glioblastoma cell lines were seeded on 10 mm coverslips and incubated at 37°C and 5% CO2 until reaching 60-70% confluence. When the cells reached the desired confluence, the culture medium was added in the presence or absence of 5 mM lactisole or vehicle control (DMSO), or 1 μM staurosporine (apoptosis positive control), followed by incubation for 48 hours. After incubation, CellEvent^TM^ (5 μM) and propidium iodide (1:1000) were added and the cells were incubated at 37°C and 5% CO 2 for 30 minutes. The culture medium was discarded, the cells were washed with PBS 1x and fixed with 4% PFA for 10 minutes, followed by a 10-minute incubation at room temperature with the Hoechst 33342 to nuclei staining. After washing, cells were mounted onto microscope slides and visualized under an Axio Imager Z2 (Carl Zeiss) microscope using a magnification of 40x (Plan-Apochromat 40x/1.3 Oil DIC M27).

### 2.4 Analysis of the role of STR on the cancer cells’ metabolism

#### 2.4.1 Effects of TAS1R3 inhibition in the glucose uptake

The glucose consumption assay was performed to determine the effect of STR inhibition in the amount of glucose absorbed by tongue cancer UPCI-SCC-154, and glioblastoma U-87MG, SNB-19 and U-373MG cells, using a deoxyglucose analogue (2-NBDG) fluorescent marker. To perform this assay, the commercial Glucose Uptake Cell-Based Assay Kit was used. Briefly, the cells were seeded in black clear-bottom 96-well plates and incubated overnight at 37°C and 5% CO2. Thereafter, the culture medium was discarded, and the cells were subjected to glucose starvation, in the presence or absence of 5 mM lactisole or vehicle control. After 4 hours of incubation at 37°C and 5% CO2, 100 μg/mL of 2-NBDG was added to each well and incubated for 30 minutes. Then, the cells were washed with cell-based assay buffer and centrifuged at 400 g for 5 minutes at RT. Subsequently, a solution of Hoechst 33342 diluted 1:1000 in cell-based assay buffer was added to label the cell nuclei, followed by centrifugation at 400 g for 5 minutes at RT. The supernatant was discarded, and 100 μL cell-based assay buffer were added to each well. Finally, the fluorescence of the Hoechst 33342 was read at excitation/emission wavelengths 361/497 nm and the fluorescence of glucose 2-NBDG at 485/535 nm on a SpectraMax Gemini spectrofluorometer (Molecular Devices). Glucose uptake was given by the ratio between 2-NBDG and Hoechst 33342 fluorescence.

#### 2.4.2 Effects of TAS1R3 inhibition in L-lactate production

Culture media from the MTT and migration experiments with lactisole (described in sections 2.3.1 and 2.3.2) were collected and used for the determination of extracellular L-lactate, following the manufacturer’s instructions. This assay allowed the detection of L-Lactate in a fluorescence-based method, in which lactate dehydrogenase catalyses the oxidation of L-lactate to pyruvate, with a consequent reduction of NAD+ to NADH, which in turn reacts with the fluorescent substrate and results in a fluorescent product. Due to the presence of lactate dehydrogenase in the samples, and to prevent the conversion of lactate to pyruvate, the samples were deproteinized with 0.5 M metaphosphoric acid and centrifuged at 10 000 g during 5 minutes at 4°C. Then, the supernatant was discarded and 50 μL of potassium carbonate were added to maintain the pH. The samples were centrifuged again under the same conditions, the supernatant was collected, and samples were diluted 1:2 in an assay buffer solution. In an opaque black 96-well plate, 20 μL of L-lactate standard samples with known concentrations (0-1000 μM) and 20 μL of the samples of interest were added in duplicate. Then, 100 μL of buffer solution, 20 μL of cofactor mix and 20 μL of fluorescent substrate were added to each well. Reactions were initiated by the addition of 40 μL of enzyme mix, and the plates were incubated, protected from light, for 20 minutes at RT. Fluorescence values were read in a SpectraMax Gemini spectrofluorometer at 530-540 nm excitation and 585-595 nm emission, and the concentration of L-Lactate was calculated as recommended by the manufacturer.

### 2.5 Evaluation of the impact of TAS1R3 inhibition in glioblastoma spheroids

The human glioblastoma cell lines (U-87MG, SNB-19 and U-373MG) were used for spheroid formation. Cells were seeded at varying densities (2.5×10^3^ U-87MG and U-373MG cells/well, and 5×10^3^ SNB-19 cells/well) into Nunclon™ Sphera™ 96-Well, U-bottom, low-attachment microplates (ThermoFisher Scientific). To promote uniform cell aggregation and spheroid formation, plates were centrifuged at 800 rpm for 1 minute immediately after seeding. Spheroids were allowed to form undisturbed for 48 hours at 37°C in a humidified atmosphere containing 5% CO2.

After 48 hours, spheroids were embedded in 2% GelTrex™ (Thermo Fisher Scientific), a synthetic extracellular matrix substitute, to assess invasive capacity. GelTrex™ was added directly to each well to allow 3D matrix formation, and the microplate was gently agitated on an orbital shaker for 5 minutes to ensure even distribution of the matrix. Then, spheroids were exposed either to 5 mM lactisole or vehicle control.

Brightfield images of the spheroids were acquired every 24 hours over a period of 7 days using a Zeiss Axio Observer Z1 inverted microscope equipped with an A-Plan 5x/0.12 Ph0 M27 objective. Spheroid diameters were quantified by image thresholding using Fiji (ImageJ) software and compared at each time point to the initial diameter at 0 hours.

### 2.6 Data Analysis

Statistical analysis and dataset comparisons were performed using GraphPad Prism 9.0

(GraphPad Software). Statistical significance between two or more groups was determined by Unpaired t test or ANOVA followed by the software’s recommended multiple comparisons post-hoc test, respectively. Results are presented as mean ± SEM of at least three independent experiments, and data were considered statistically different for a p-value < 0.05.

## 3. Results

### 3.1 TAS1R3 inhibition impairs the proliferative capacity but does not trigger cell death in tongue cancer and glioblastoma cells

The effects of TAS1R3 inhibition with lactisole in the viability and migration were assessed in the tongue cancer UPCI-SCC-154 cells, and in three glioblastoma cell lines U-87MG, SNB-19 and U-373MG (Figures 1 and 2).

**Figure 1-.**
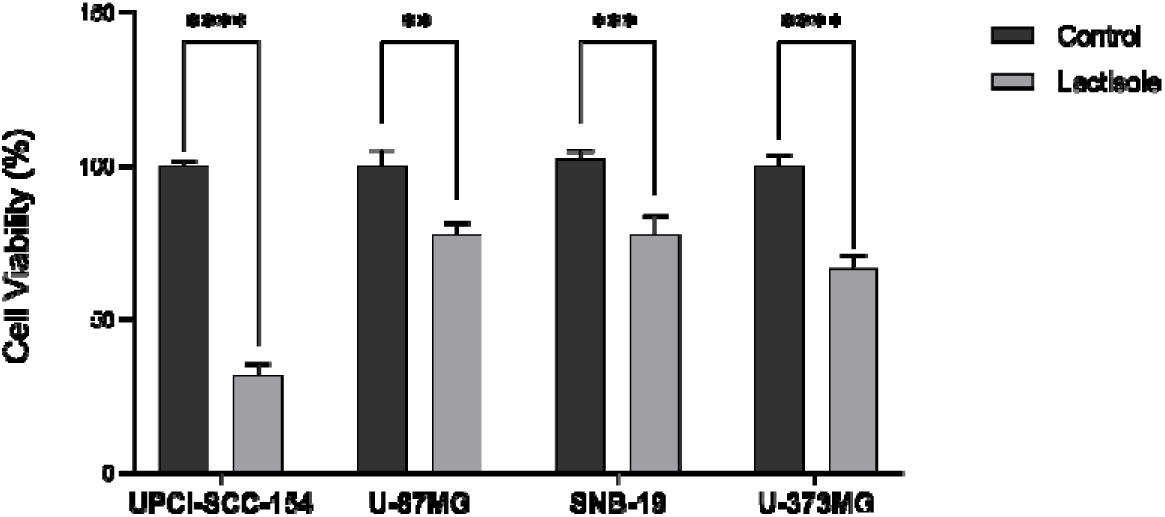
Effect of TAS1R3 inhibition with lactisole on tongue cancer and glioblastoma cell lines’ viability. Effect of TAS1R3 inhibition in the viability of tongue cancer UPCI-SCC-154 cells, and glioblastoma U-87MG, SNB-19 and U-373MG cells grown in culture media containing DMSO (vehicle control), in the presence or absence of 5 mM lactisole, for 48 hours. Bar graphs represent mean ± SEM. Statistical analysis was performed by unpaired t test. [N≥3 independent experiments; **p<0.01, ***p<0.001 and ****p<0.0001].

**Figure 2-.**
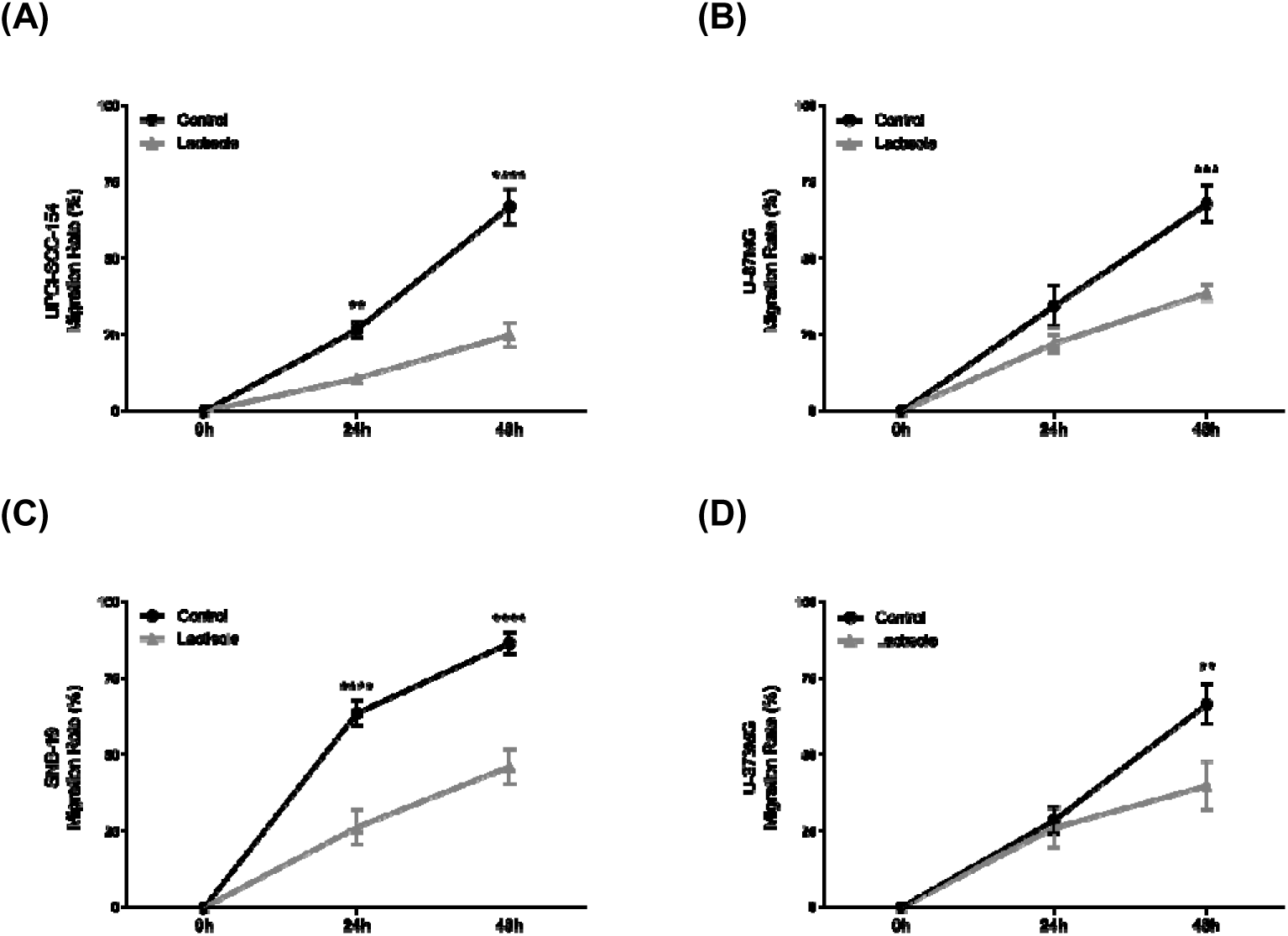
Effect of TAS1R3 inhibition with lactisole on tongue cancer and glioblastoma cell lines’ migration rate. Effect of TAS1R3 inhibition in the migration rate of tongue cancer UPCI-SCC-154 cells, and glioblastoma U-87MG, SNB-19 and U-373MG cells grown in culture media containing DMSO (vehicle control), in the presence or absence of 5 mM lactisole, for 48 hours. Bar graphs represent mean ± SEM. Statistical analysis was performed by two-way ANOVA followed by Sídák’s multiple comparisons test. [N≥3 independent experiments; **p<0.01, ***p<0.001 and ****p<0.0001].

We found that TAS1R3 inhibition reduced the viability of UPCI-SCC-154 cells from tongue cancer (68.33%±2.43), and U-87MG (22.48%±1.32), SNB-19 (24.35%±3.43) and U-373MG (33.60%±1.25) glioblastoma cells (Figure 1).

Then, we evaluated the effects of TAS1R3 inhibition with lactisole on the migratory capacity of tongue cancer and glioblastoma cells (Figure 2). Overall, we observed that the migration and proliferation rates of the control group were higher when compared to the group of cells where the STR was inhibited. The migration rate of U-87MG and U-373MG cells (Figures 2B and 2D) was statistically different between the control and lactisole groups only after 48 hours of incubation (67-68% *versus* 39-40%). On the other hand, the TAS1R3 inhibition was shown to be effective in decreasing the migration ability of UPCI-SCC-154 and SNB-19 cells (Figure 2A and 2C) after 24 and 48 hours of incubation with lactisole (27-64% *versus* 11-26% and 67-87% *versus* 25-46%, respectively).

Our observations put in evidence that STR inhibition, via blockage of the TAS1R3 subunit, impairs the proliferative and migration capacity of glioblastoma and tongue cancer cells. To elucidate the mechanism by which STR inhibition leads to decreased proliferation and migration of glioblastoma cells, we proceeded with the assessment of cell death by staining apoptotic or necrotic cells (Figure 3). Interestingly, neither apoptosis nor necrosis were observed, suggesting that lactisole does not trigger cell death in glioblastoma cells.

**Figure 3-.**
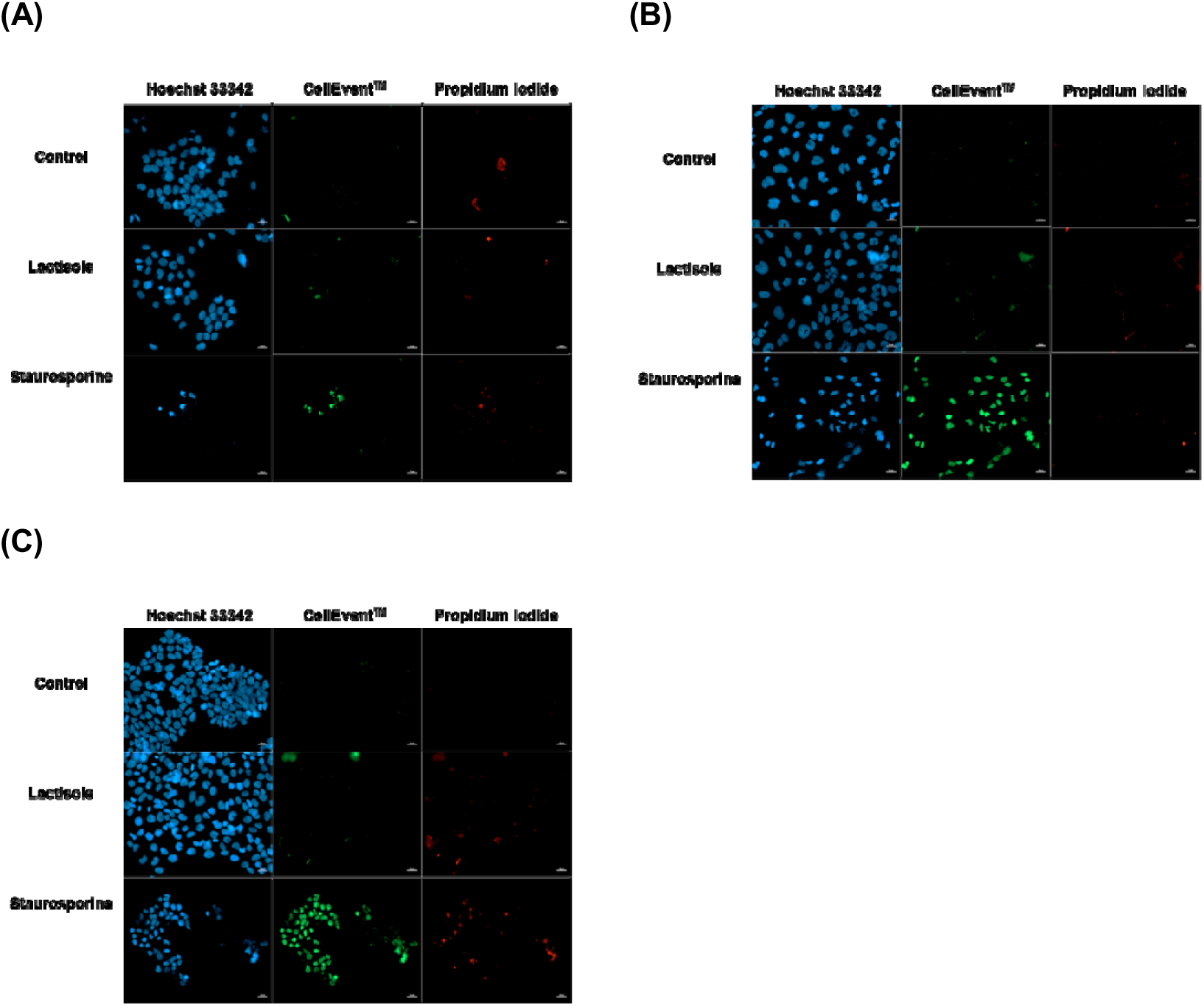
TAS1R3 inhibition did not induced cell death in glioblastoma cells. Cell death assessment was performed by staining apoptotic (CellEvent^TM^; green) or necrotic (propidium iodide; red) glioblastoma cells. (A) U-87MG, (B) SNB-19 and (C) U-373MG grown in culture media containing DMSO (vehicle control), in the presence or absence of 5 mM lactisole, for 48 hours. A positive control for apoptosis was used by incubating the cells with 1 µM staurosporine for 48 hours. Nuclei were stained with Hoechst 33342 (blue). Scale bar: 20 µm.

### 3.2 TAS1R3 inhibition impairs glucose metabolism in tongue cancer and glioblastoma cells

The effect of TAS1R3 inhibition on glucose uptake by tongue cancer and glioblastoma cells, was assessed using a fluorometric glucose uptake assay was performed using the 2-NBDG marker. As depicted from figure 4, blocking TAS1R3 reduced the glucose uptake of tongue cancer UPCI-SCC-154 cells by 30%, and U-87MG and SNB-19 glioblastoma cells by 28% and 64%, respectively. No differences were observed in the glioblastoma U-373MG cell line (Figure 4).

**Figure 4-.**
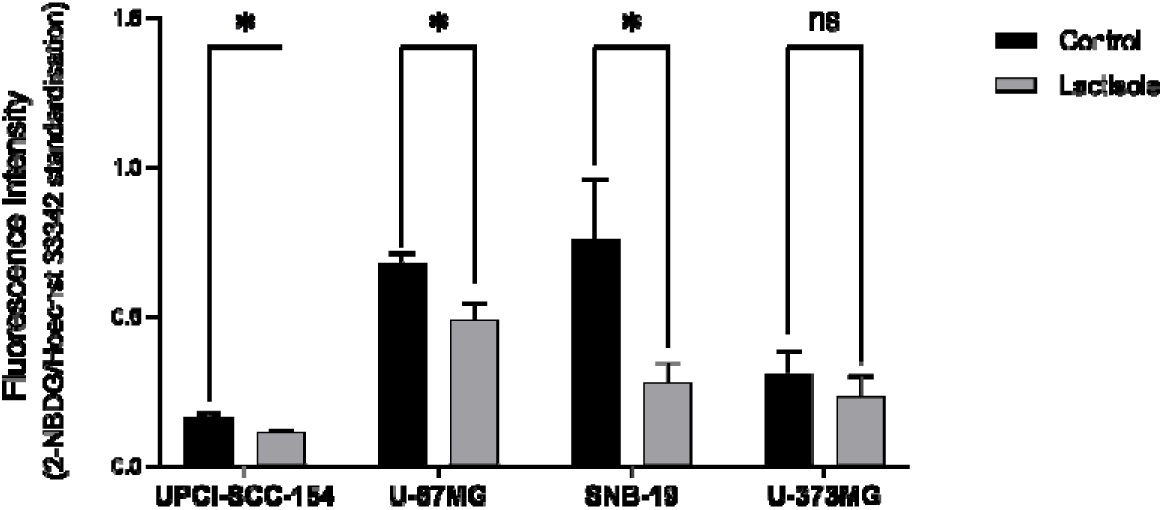
Effect of TAS1R3 inhibition in the glucose consumption by tongue cancer and glioblastoma cells. Effect of TAS1R3 inhibition in the glucose uptake by UPCI-SCC-154 tongue cancer cells, and U-87MG, SNB-19 and U-373MG glioblastoma cells glucose-starved during 4 hours, in the presence or absence of 5 mM lactisole or vehicle. Bar graphs represent mean ± SEM. Statistical analysis was performed by unpaired t test. [N≥3 independent experiments; *p<0.05, ^ns^non-significant].

Considering that many cancer cells rely on glycolysis to proliferate, we aimed at investigating the effect of TAS1R3 inhibition in L-lactate production. In line with the results obtained for the glucose uptake, a decrease between 14-28% in the concentration of extracellular L-lactate produced by the UPCI-SCC-154 tongue cancer, and U-87MG and SNB-19 glioblastoma cells (Figure 5) was also observed.

**Figure 5-.**
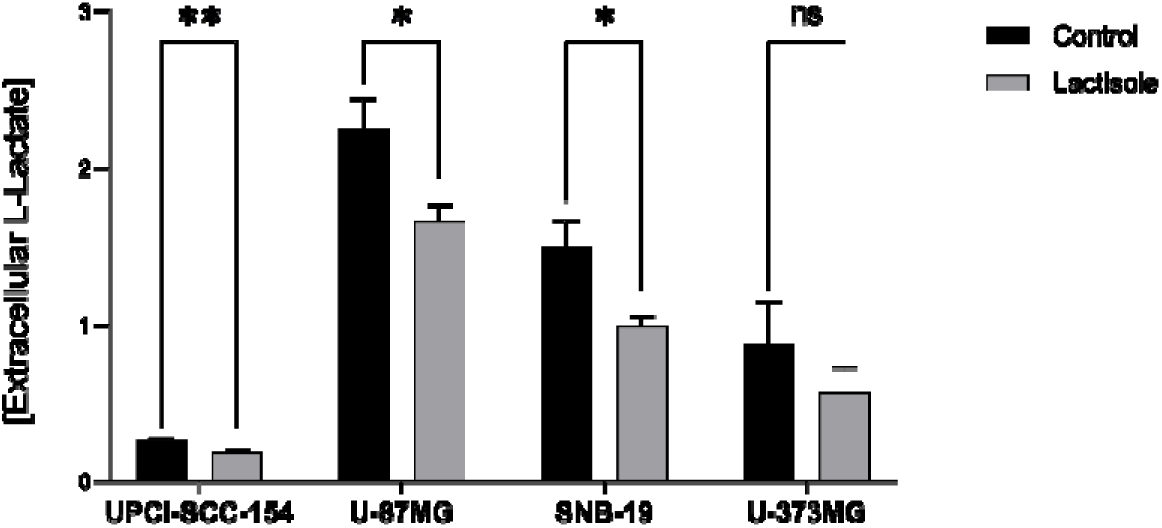
Effect of TAS1R3 inhibition in the L-lactate production by tongue cancer and glioblastoma cells. Effect of TAS1R3 inhibition in the concentration of extracellular L-lactate produced by the UPCI-SCC-154 tongue cancer cells, and U-87MG, SNB-19 and U-373MG glioblastoma cells grown in culture media containing DMSO (vehicle control), in the presence or absence of 5 mM lactisole, for 48 hours. Bar graphs represent mean ± SEM. Statistical analysis was performed by unpaired t test. [N≥3 independent experiments; *p<0.05, ^ns^non-significant].

### 3.3 TAS1R3 inhibition affects the invasion rate of glioblastoma spheroids

The next step was to investigate the anticancer effects of STR inhibition by lactisole in a more physiologically relevant model. Thus, we examined the effects of lactisole in glioblastoma cell lines spheroids, a pre-clinical model that better mimic the 3D architecture of tumours.

Invasion assays using GelTrex™-embedded glioblastoma spheroids revealed that STR inhibition with lactisole significantly reduced the invasion of U-87MG cells (Figure 6A), while no effect was observed in SNB-19 and U-373MG cells (Figures 6B and 6C).

**Figure 6-.**
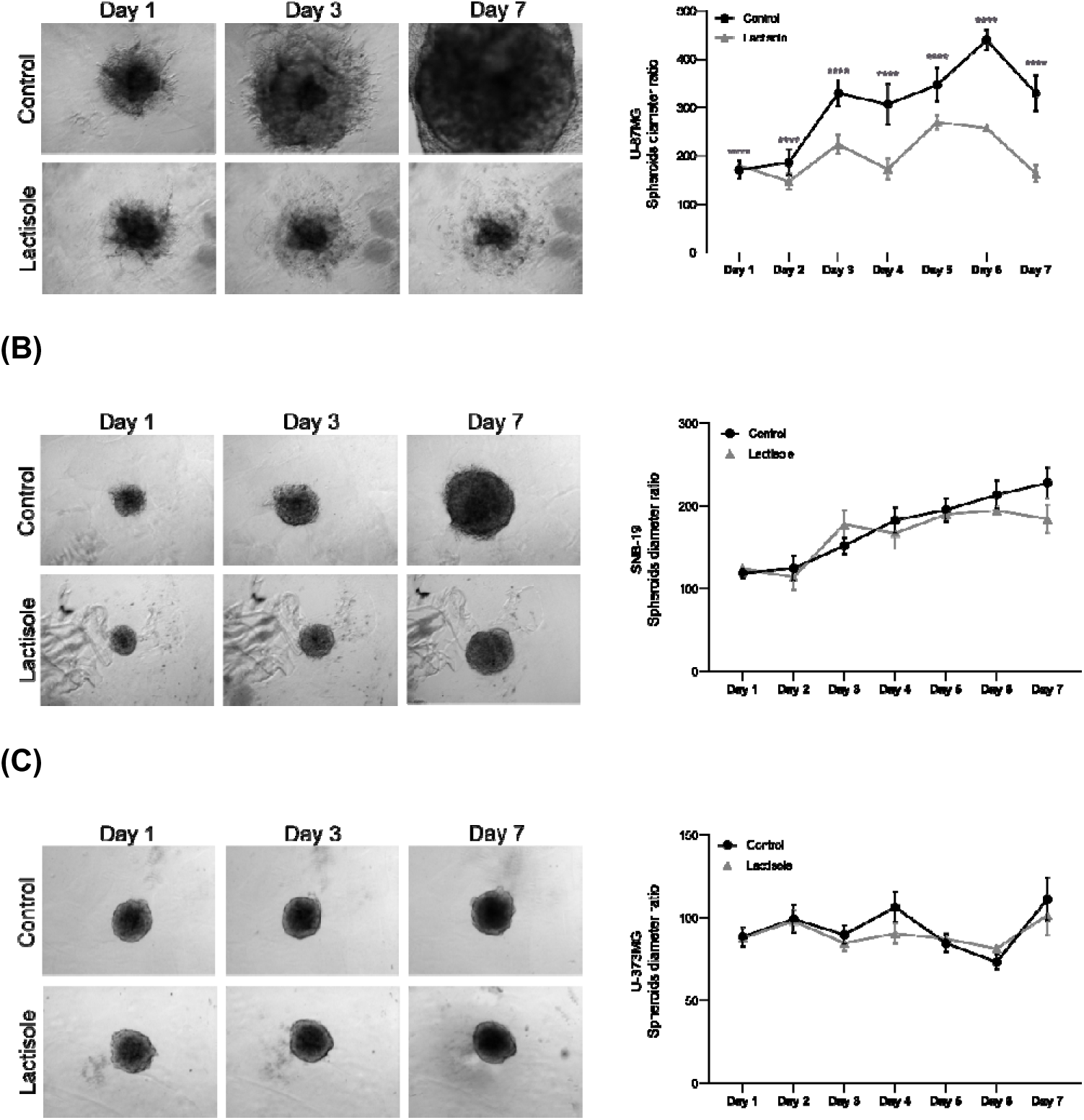
TAS1R3 inhibition reduces invasion of U-87MG glioblastoma spheroids. Invasion assays were performed by embedding glioblastoma spheroids in 2% GelTrex™, in the presence or absence of either 5 mM lactisole or vehicle control (DMSO). Spheroids of (A) U-87MG, (B) SNB-19, and (C) U-373MG were imaged every 24h for 7 days using brightfield microscopy, and the invasion was assessed by measuring the spheroids diameter. Scale bar: 200 µm.

## 4. Discussion

Glioblastoma is a highly aggressive tumour with increased proliferative and invasive capacity, and is markedly dependent on the metabolic process of aerobic glycolysis, also known as the Warburg effect, which relies on sustained glucose availability (Liberti & Locasale 2016; Linkous & Yazlovitskaya 2011; Schröder et al. 1991). Given the critical role of glucose metabolism, our experiments aimed at assessing whether STR could act as a sensor of glucose availability and a regulator of glucose metabolism in glioblastoma, as reported in several organs and tissues. For example, in the intestine, STR signalling increases the rate of glucose absorption by regulating a glucose transporter in enterocytes (Smith et al 2018). The administration of non-caloric sweeteners in a gastric cancer cell line was shown to induce serotonin secretion dependent on the activation of the TAS1R3 subunit (Zopun et al. 2018). In wild-type rats, a sucrose-rich diet reduced the glucose uptake by rapidly inducing STR downregulation in the gut (Smith et al. 2018). In another study with rat brains, type 1 taste receptors and G protein-associated genes, are expressed in different nutrient-sensitive regions of the forebrain, including the hypothalamus, hippocampus, habenula, cortex, and in the choroid plexus (Ren et al. 2009). Thus, G protein-coupled taste receptors may serve as membrane-bound chemosensors, with STR being a major candidate for brain glucosensing. A recent study carried out in astrocytes and in neurons from several nutrient-sensitive regions of the midbrain, suggests that STR has a role in controlling glucose metabolism. In addition, STR expression was detected in tanycytes specialized glial cells of the third brain ventricle, a brain region highly sensitive to glucose (Ren et al. 2009; Welcome & Mastorakis 2018). In the hypothalamus, expression levels of taste-related genes appear to depend on nutritional status (Ren et al. 2009), suggesting that taste signalling mechanisms in the brain could be involved in the assessment and regulation of nutrient uptake, particularly glucose.

In a previous study, we observed differential TAS1R2 and TAS1R3 protein expression levels between the glioblastoma cell lines analysed (Costa et al. 2025 - bioRxiv submission ID BIORXIV/2025/661264). In cancer cells, including glioblastoma, higher STR expression is likely to be advantageous for tumour progression since it could enhance glucose sensing and uptake from the tumour microenvironment (Jiang 2017). As seen in other types of cancer, glioblastoma growth and chemoresistance is highly enhanced by the metabolic switch to aerobic glycolysis, which favours the lactate release to tumour microenvironment and the weakening of the immune response in the tumour microenvironment (Denko 2008; Desbats et al. 2020). Thus, we investigated the effect of TAS1R3 subunit inhibition, with lactisole, in the proliferative capacity, migration, and apoptosis of the three glioblastoma cell lines. Interestingly, STR inhibition consistently reduced the viability and migration in three glioblastoma cell lines and in the tongue cancer UPCI-SCC-154 cell line, used as a positive control for STR expression. The reduction in viability and migration was particularly marked in UPCI-SCC-154 tongue cancer cells. This may be due to the higher STR expression levels expected in these cells. In the three glioblastoma cell lines, both viability and migration were significantly impaired, albeit with distinct kinetics. Interestingly, neither apoptosis nor necrosis was observed in the glioblastoma cells, suggesting that the reduced proliferation might result from metabolic stress or growth arrest, rather than cell death, as a consequence of reduced glucose uptake. This is consistent with previous findings showing that glucose deprivation induces senescence and cell cycle arrest in cancer cells and astrocytes (Han et al. 2015; Gao et al. 2022). In endometrial cancer cells exposed to low glucose concentrations, the migratory capacity was reduced, and the cell growth and proliferation were inhibited, indicating that the antiproliferative effects exerted by glucose deprivation could be attributed to cell cycle arrest (senescence) (Han et al. 2015). Recently, it was shown that astrocytes subjected to low glucose conditions underwent senescence (Gao et al. 2022).

As described before, cancer cells adapt to the low energy yield of glycolysis by increasing glucose uptake to support the high glycolytic rate (Liberti & Locasale 2016). This can benefit cancer cells by providing both ATP and key metabolic intermediates necessary for rapid cell proliferation but depends on high glucose concentrations (Liberti & Locasale 2016; Lunt & Vander Heiden 2011). To further investigate the metabolic impact of TAS1R3 inhibition, we assessed glucose uptake and L-lactate production. Lactisole treatment significantly decreased glucose uptake and lactate secretion in UPCI-SCC-154 tongue cancer cells, and in U-87MG and SNB-19, but not in U-373MG glioblastoma cells, suggesting cell-line-specific metabolic responses. In fact, U-373MG represent tumour core-like cells, which are less exposed to oxygen and circulating nutrients, including glucose (Diao et al. 2019; Vijayanathan & Ho 2025). These results reinforce the idea that STR activity contributes to glycolytic metabolism in glioblastoma. Similar results were observed in other cancer models, where blocking STR or glycolysis reduced lactate production and proliferation (Andrade et al. 2018; Sun et al. 2017).

Importantly, to address the functional relevance of STR inhibition in a more physiologically relevant model, we evaluated glioblastoma spheroids embedded in an extracellular matrix. Lactisole treatment reduced the invasive capacity of U-87MG spheroids, although no significant effect was observed on SNB-19 or U-373MG spheroids. This may be attributed to the fact that U-87MG cells exhibit collective invasion patterns and resemble cells from the invasive tumour margin, areas more associated with recurrence and therapeutic resistance (Andrieux et al. 2023; Diao et al. 2019).

In summary, we demonstrate that STR plays an important role in glioblastoma cell metabolism by acting as a glucosensor of glucose availability. Our findings show that TAS1R3 inhibition affects glioblastoma cell viability, migration, and metabolism, particularly in cell lines with higher sensitivity to metabolic disruption. In line with this, STR appears to be involved in sustaining glioblastoma cell proliferation and invasion, likely by regulating glucose uptake and glycolysis. Because STR inhibition with lactisole did not induce cytotoxicity, it may offer a therapeutic window to impair tumour progression with minimal or no negative side effects. Notably, STR blockade has already been shown to exert antidiabetic effects in humans, without reported adverse effects (Teff et al. 2010). These are promising features for targeting this receptor as an anticancer therapeutic strategy.

## Conflicts of interest

The authors declare that they have no known competing financial interests or personal relationships that could have appeared to influence the work reported in this paper.

## Acknowledgements

This work was supported by the Fundação para a Ciência e Tecnologia (FCT, Portugal) project grants (PTDC/BIM-ONC/7121/2014, UID/Multi/00709/2013 and UID/Multi/00709/2019), by CENTRO 2020 and Lisboa 2020 project grant (POCI-01-0145-FEDER-016822), and FEDER funds through the POCI – COMPETE 2020 – Operational Programme Competitiveness and Internationalization in Axis I – Strengthening research, technological development and innovation (POCI-01-0145-FEDER-007491). The work was also supported by the Portuguese Cancer League, through the Bolsa de Investigação em Onocologia Dr. Rocha Alves 2022. Ana R. Costa was recipient of a PhD fellowship (UI/BD/151025/2021) funded by FCT through the Portuguese state and EU budgets through the European Social Fund. Ana C. Duarte was recipient of a grant from CENTRO 2020 program through the ICON project (Interdisciplinary Challenges On Neurodegeneration; CENTRO-01-0145-FEDER-000013). We also acknowledge the support of the Portuguese Platform of Bioimaging (PPBI) [PPBI-POCI-01-0145-FEDER-022122] and the resources provided by the Fluorescence Microscopy Unit of RISE-Health, UBI.

## Notes

### Competing Interest Statement

The authors have declared no competing interest.

